# Motor learning and adaptation in bird flight

**DOI:** 10.64898/2026.01.20.700397

**Authors:** Clementine Bodin, Jasmin C.M. Wong, Shane Windsor, Sarah C. Woolley

**Affiliations:** Department of Biology, McGill University; School of Civil, Aerospace and Design Engineering, University of Bristol; Center for Research on Brain, Language, and Music

## Abstract

Birds are capable of performing elaborate flight maneuvers in variable environmental conditions. While flight is an adaptable and skilled motor behavior, we know surprisingly little about how birds master this ability. Across species, skilled motor behaviors show practice-related changes or improvements in performance and understanding which features of a motor behavior change with learning can lend insight into the constraints, flexibility, and optimization of motor behavior. Here, we combined high-speed video recordings and pose-tracking software to analyze the kinematics of thousands of flights in zebra finches over multiple days of flight training and over different distances. Small birds, such as the zebra finch, use an intermittent ‘flap-bounding’ flight style that alternates between flapping phases and flexed-wing bounding phases. We found that birds increase their flight speed and the time spent bounding and reduce variability in the bound position over ten days of flight performance. These motor changes to flight show savings, as performance is maintained even after two months without flight experience. Moreover, these same parameters are adjusted when birds fly longer distances, indicating that they may be key to flight flexibility. We built kinematic models to determine what features birds might be optimizing toward with learning and found that the data was best fit by a model simultaneously optimizing for minimum energy and flight duration. Taken together, our data highlight that flap-bounding flight shows hallmarks of skilled motor learning and lend new insight into the function of bounds.

## INTRODUCTION

Across species, motor systems can produce a tremendous diversity of behaviors, and variation in motor performance is often under natural selection, sexual selection, or both. A hallmark of skilled motor behaviors is that they show practice-related changes or improvements in performance^1–3^. Just as practice in humans improves our ability to play music or ride a bike, other animals exhibit motor behaviors that require practice for optimization, for example vocal learning in songbirds ^4–8^ and skilled reaching and grasping in rodents and primates ^9–11^. Understanding which features of a motor behavior change with learning can lend insight into the constraints, flexibility, and optimization of motor behavior.

Birds are capable of elaborate flight acrobatics, which they can use not only for locomotion but also to evade predators, hunt prey, or perform courtship displays. Moreover, birds can adjust flight based on environmental conditions, for example changes to wind speed^12^ and obstacles^13,14^, indicating flight is a challenging and flexible motor skill. Birds display a wide range of flight styles, from the soaring of vultures to the hovering of hummingbirds, shaped by environmental pressures and anatomical constraints^15–18^. Small birds, such as the zebra finch, use an intermittent ‘flap-bounding’ flight style that alternates between flapping phases and flexed-wing bounding phases^18^. Multiple studies have modelled flap-bounding flight to understand the function and evolutionary significance of this flight style. In particular, it has been alternately suggested that flap-bounding flight may be less energy-intensive than continuous flapping or that intermittent bounds may be a way to maximize horizontal velocity or to adjust flight altitude or speed^12^. While studies modelling flap-bounding flight lend insight into the kinematics and biomechanics of flight behavior, there is a notable lack of studies on motor learning and practice in flight.

Here, we developed a small, adjustable flight chamber and recorded flight performance over many days of flight experience as well as over multiple distances to investigate whether flap-bounding flight exhibits hallmarks of motor practice, including changes to the speed and variability of flight. We uncovered several features that change with flight experience and with distance, indicating that these features may be key to motor flexibility of flight. In addition, we hypothesized that investigation of flight learning and practice could elucidate the optimization of the flap-bounding flight style. To this end, we used the flight kinematics and trajectories of birds given extensive flight practice to generate kinematic models containing different optimization parameters to gain a better understanding of the function of flap-bounding flight. Taken together, our data indicate that flap-bounding flight is a flexible motor skill that improves with practice. The learning-induced changes to flight kinematics lend needed insight into the function of the flap-bounding flight style.

## RESULTS

Flight is a highly adaptable and flexible motor behavior. However, we have little understanding of the degree to which flight changes with experience or what parameters of flight are adjusted depending on flight distance. Here, we used markerless tracking of zebra finches during flight (DeepLabCut, see Methods, Fig. 1) to analyze the movement of the beak, wingtips, and tail in time and space. We compared these kinematic data within the same cohort of birds over ten days of flight experience and with increasing flight distance.

**Figure 1.**
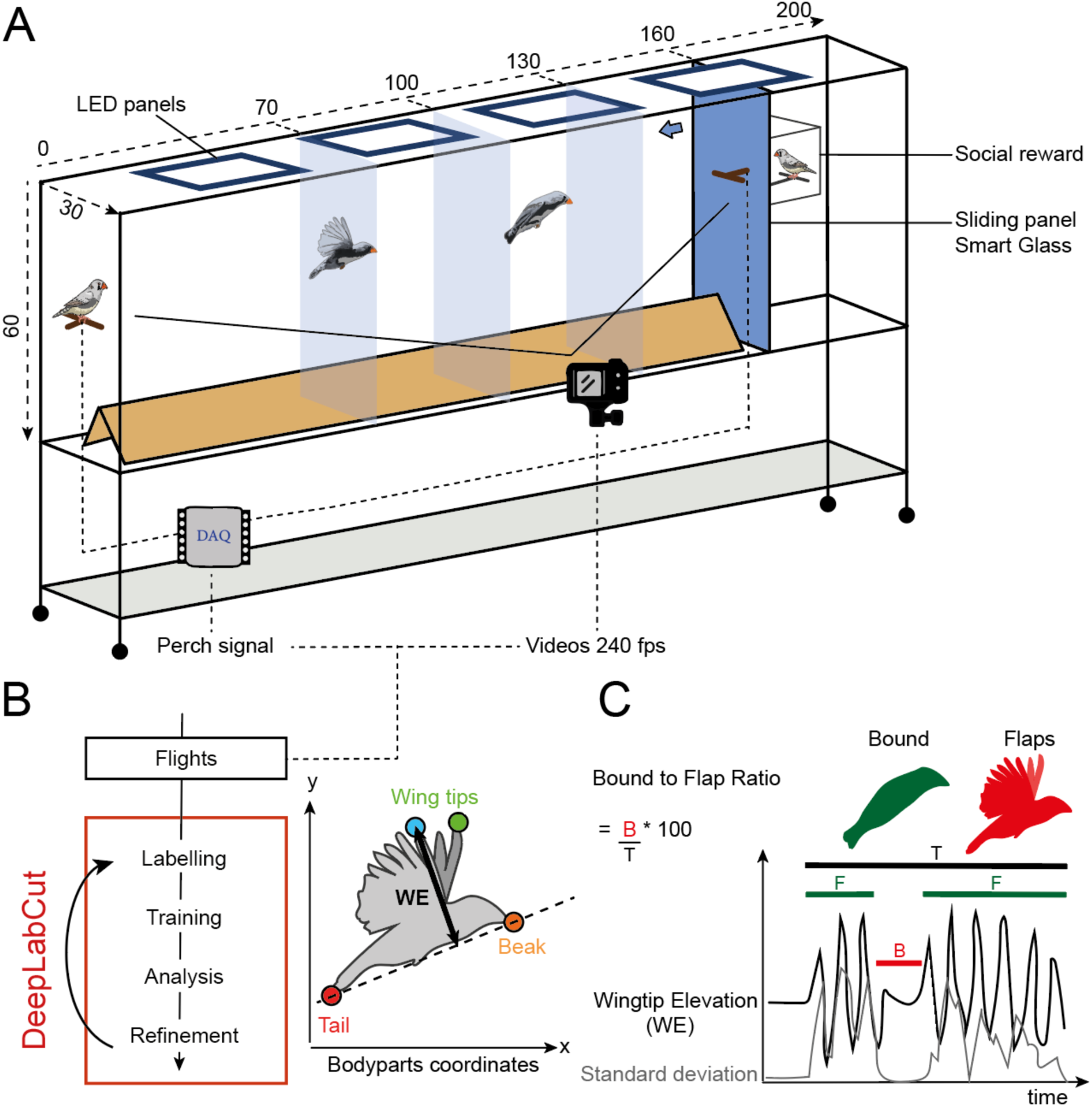
(A) Diagram of adjustable flight chamber. Birds flew along a plexiglass corridor (70 to 160 cm long). Perches at each end connected to limit switches to record take-off and landing times that could be aligned with videos (240 fps). Landing on the perch on the right side triggered a piece of smart glass to become clear and allow the bird to view another bird (social reward). (B) Analysis pipeline. Recorded flights were extracted based on perch movement times and the wingtips, beak, and tail were labelled using AI-based pose-tracking software (DeepLabCut) to determine x and y coordinate positions over time. Elevation of the wingtips relative to the axis created by the beak and tail (B) was used to identify epochs of bounding and flapping (C).

### Flight kinematics change with experience

We first investigated the flexibility of flight by quantifying the degree to which flight kinematics change with experience. We recorded birds daily for ten days as they flew over a fixed flight distance (70cm, Fig. 1A). Zebra finches alternate between flapping and bounding flight phases where bounds are periods without flapping when birds bring their flexed wings in close to the body. Our analysis of wingtip elevation allowed us to distinguish between flapping and bounding phases and to calculate detailed features of flight (see Methods; Fig. 1C). Overall, birds adjusted their flight speed as well as the proportion of the flight they spent bounding over the course of the ten days of experience. For example, Fig 2A shows data from the initial, middle, and final flights for a single bird who increased her flight speed from 1.31 to 1.60 m/s. This bird also introduced a bound at the middle of the flight corridor. To quantify changes to bounding across flights and between birds we calculated a bound ratio (BR) as the proportion of the total flight spent in a bound. In this example the BR increases from 0% (no bounds) on Day 1 to 35.1% of the flight duration on Day 10.

**Figure 2.**
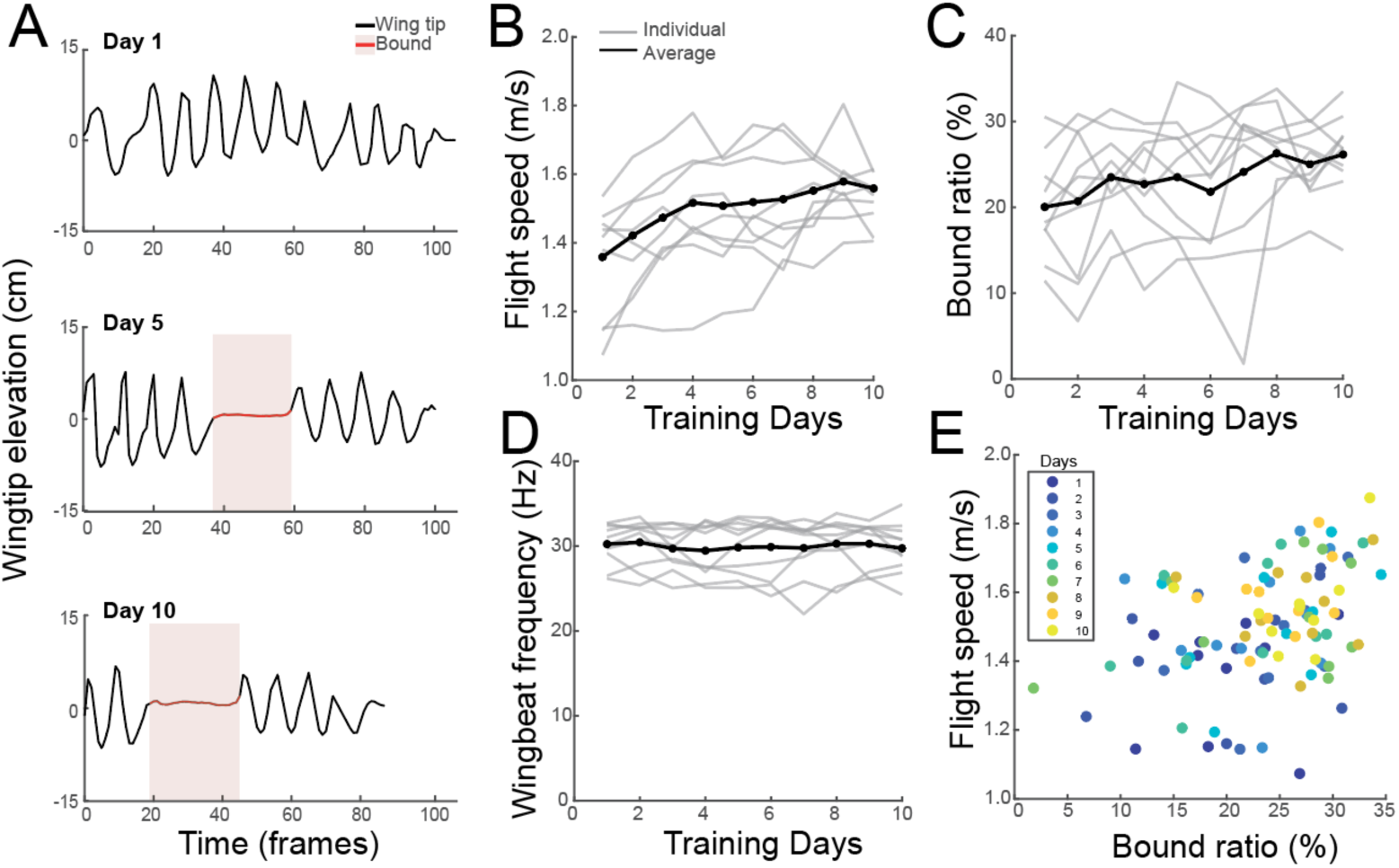
(A) Wingtip elevation of a single bird on three example flights during the 10-day training highlighting the increase in speed, as indicated by shorter time to complete the flight, and an increase in bounding with no bound on Day 1 (top panel), a short bound on Day 5 (middle panel), and a long bound on Day 10 (bottom panel). Bounds are shaded in pink. Red lines indicate periods with no deviation in wingtip elevation which is used to categorize bounding. Across all individuals speed (B) and the percent of the flight spent bounding (Bound Ratio; C) increased significantly with training. (D) In contrast, wingbeat frequency remained steady. (E) Bound ratio and speed increase simultaneously. Day of training is indicated by color. Blue colors are early in training, yellow colors are later in training.

We saw similar increases in both flight speed and bounding across birds. Flight speed significantly increased with flight experience in all birds both within a day (Fig. 2B; flight daily rank, β = 0.003, p = 0.013) as well as over the ten days of flight experience (flight total rank, β = 0.002, p < 0.0001). The number of bounds for all birds among the 2,009 flights ranged from 0 to (mean = 1.25). The average BR also increased slightly but significantly over the ten days (Fig. 2C, mean BR = 23.4 ± 5.3% across bird IDs), with 8 of the 10 birds exhibiting a higher BR on Day 10 compared to Day 1 of training. Similar to flight speed, the BR increased significantly both within a day (flight daily rank, β = 0.006, p < 0.0001) and over days (flight total rank, β = 0.363, p < 0.0001), with no difference between the flight directions (β = 0.007, p = 0.410). In contrast, the wingbeat frequency, measured as the number of flaps per second, was not affected by either the total or daily rank of the flight (β = 0.24 and 0.88 respectively; p>0.10 for both). Moreover, there was little individual variation in wingbeat frequency either within or across days (Fig. 2D). Finally, we found that within our flight speed range of 0.62 to 2.28 m/s, speed did not correlate with wingbeat frequency (β = -0.00, p = 0.69).

Given the increase in both flight speed and bound ratio with flight experience, we explored the degree to which these parameters were correlated. Averaging their respective daily values revealed that higher speeds were associated with elevated bound ratios, peaking notably during the final days of training (Fig. 2E). Using a mixed-effects model we analyzed the relationship between speed and BR with a possible interaction with the day of practice (see Methods). We found that speed significantly increased with BR (β = 0.006, p < 0.0001) and day of training (β = 0.015, p < 0.0001). However, while both speed and bounding increase with flight experience, the relationship between them does not vary over the ten days, indicated by the lack of an interaction between speed, bound ratio, and day (β = 5.32e-05, p = 0.708). Taken together, we find that flight experience is associated with changes in flap-bounding behavior as flight kinematics, including speed and bounding, but not wingbeat frequency, are adjusted over progressive flights.

### Birds modify similar flight parameters with experience and distance

To gain further insight into the specific features of flight that birds adjust, we tested birds on flights over different lengths of the flight chamber. As with flight training, birds increased their flight speed and altered the duration and number of bounds over different distances. For example, as she flew over increasing distances (0.7, 1, 1.3, and 1.6m) the bird in Figure 3A increased flight speed from 1.48 to 2.35 m/s. Across all birds, flight speed significantly increased with flight distance (Fig. 3B), from an average of 1.48 m/s at 0.7m to 2.19 m/s at 1.6m (β = 0.56, p < 0.0001).

**Figure 3.**
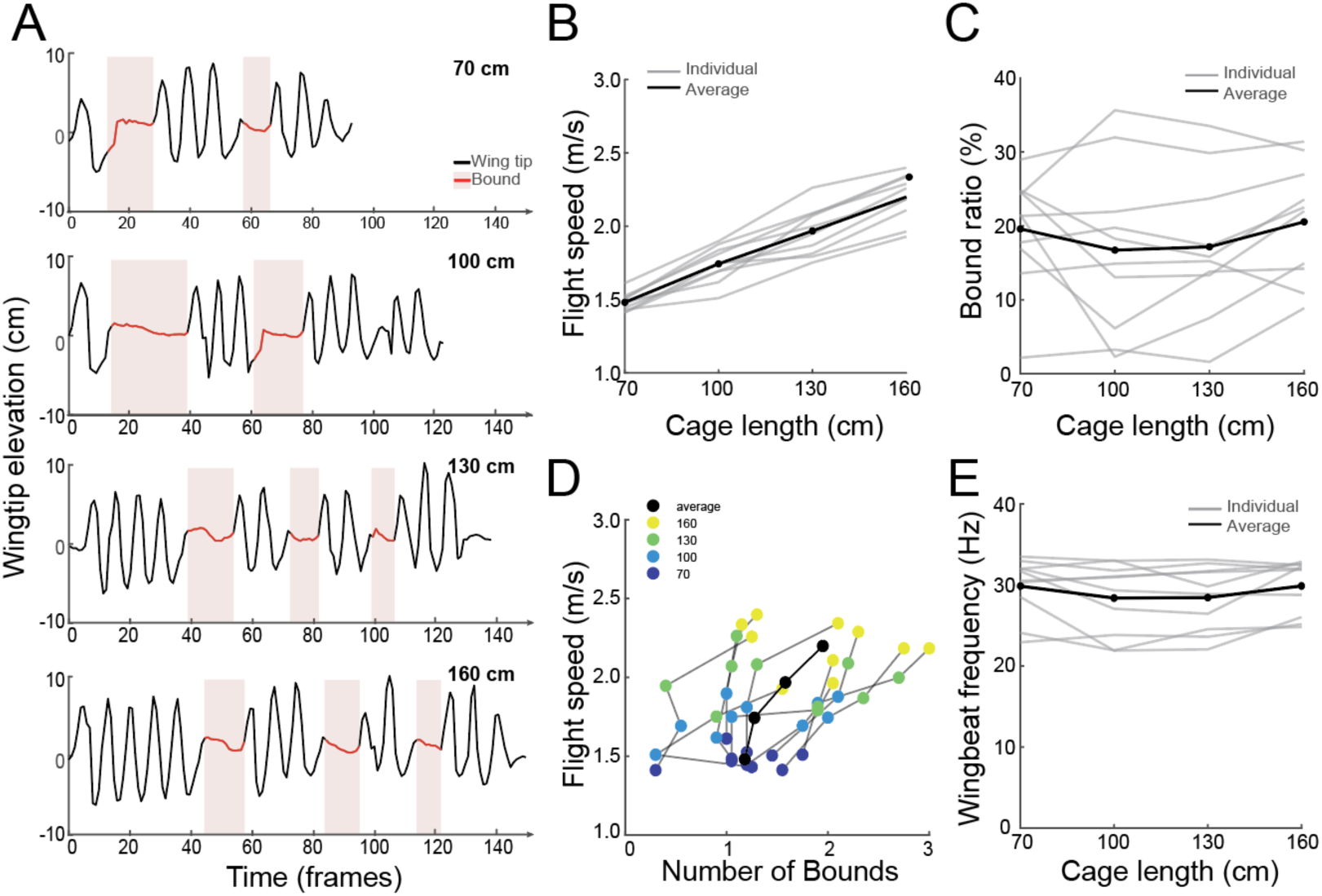
(A) Wingtip elevation of a single bird on four example flights at increasing distances, from 70 cm (top panel) to 160 cm (bottom panel), highlighting the increase in the number of bounds (pink shading, red line). Birds significantly increased flight speed with longer cage lengths (B) but did not increase the percent of time bounding (bound ratio, C). (D) Both the number of bounds and speed increase with longer cage lengths, indicated by color (yellow is for longer cage lengths). (E) There was no difference in the wingbeat frequency across the distances.

In addition to increasing flight speed with distance flown, birds also increased the number of bounds with increasing distance. For example, the bird in Fig. 3A increased the number of bounds they performed from two to three. Notably, this bird bounded at approximately similar periods for the two shortest distances, then delayed them and added a third bound for the two longest distances. However, although there were more bounds in longer flights, they were shorter in duration, resulting in similar overall percent of time spent bounding over all distances (21-25%). Across birds, we found no significant relation between the percent of time spent bounding and distance (β = -4.5, p = 0.09; Fig. 3C). However, the number of bounds increased significantly with the distance traveled (β = 1.05 p < 0.01; Fig. 3D). Finally, just as wingbeat frequency did not vary with flight experience, we also found that wingbeat frequency was impressively stable across different flight distances (β = -0.99, p = 0.39), with average values for each of the four increasing distances of 29.8, 28.4, 28.4 and 29.9 flaps/s respectively (Fig. 3E).

### Flight trajectories and bound positions become more stereotyped with experience

Birds adjust the frequency and duration of bounds over ten days of flight experience and with increasing distance. Given the flexibility in this behavioral parameter, we investigated whether there are also changes to the trajectory of flights and the location of bounds within the flight trajectories. We extracted trajectories from the steadier motion of the beak and identified the locations of bounds within those trajectories. The trajectories (n=100) made by one bird over the ten days of flight experience (Fig. 4A) highlight features of trajectories that were consistent across birds: birds flew inverted U-shaped trajectories with bounds near an altitude peak located toward the midpoint of the flight corridor’s length. Each of these features of trajectories change with flight experience.

**Figure 4.**
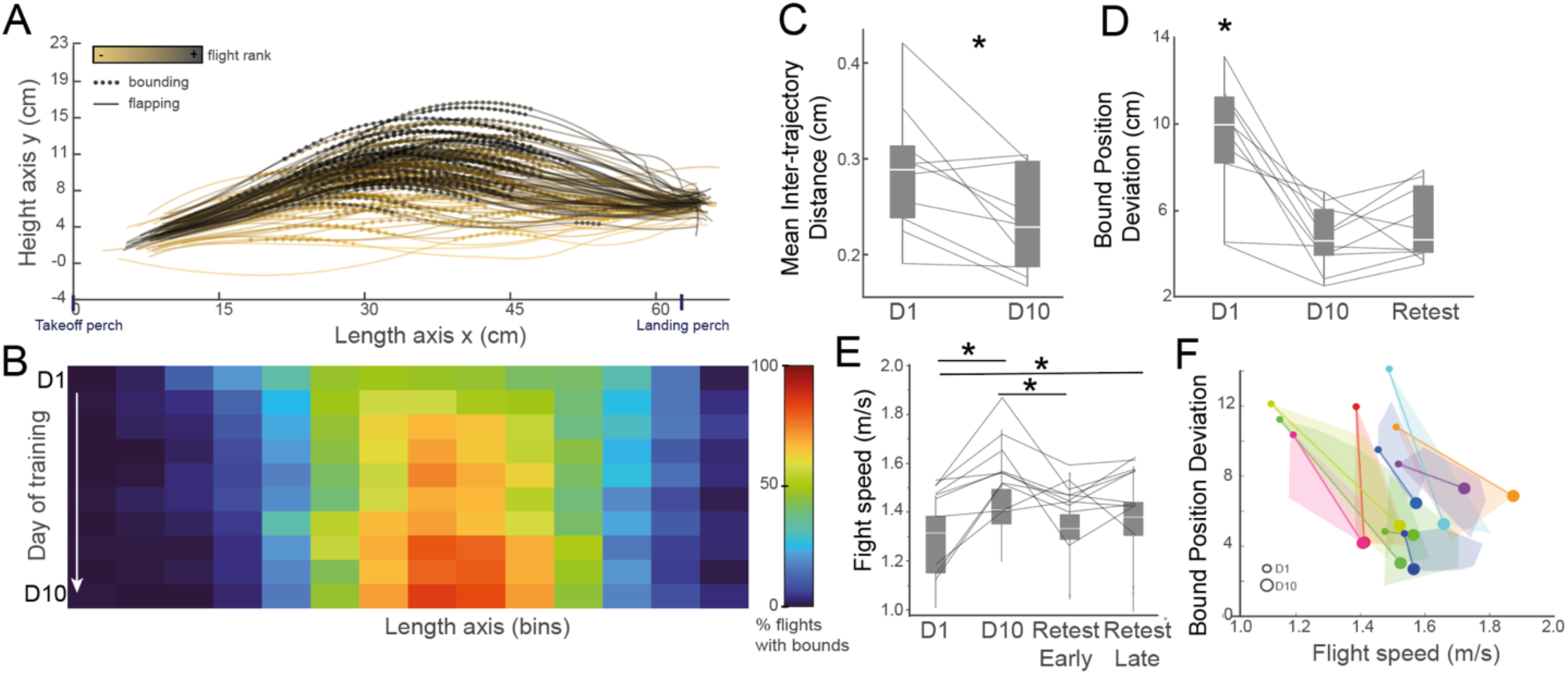
(A) Flight trajectory traces based on beak position for a single bird on all 70 cm flights. Light colors are early in training, dark colors are later in training. Diamonds indicate bounds. (B) Heat map of bound positions for all birds from Day 1 (top row) to Day 10 (bottom row). Length axis is divided into 14 bins. Color indicates the percent of flights across all birds with a bound in that bin (warmer colors are higher percent). With training, trajectory shapes became more similar, indicated by a decrease in the inter-trajectory distance (C) and a decrease in the deviation of the bound position (D). Bound position deviation remained low when birds returned to the flight chamber after two months without training (D). (E) Flight speed increases with training and recovers to trained levels within 20 flights following two months without training. (F) Speed increases and the deviation of bound position decreases simultaneously over training. Each color represents a different bird with Day 1 (small circle) and Day 10 (large circle) connected by a line.

We first looked at changes to the overall trajectory shape by comparing the Euclidean distance between trajectories performed on Day 1 with those from Day 10. The inter-trajectory distance was significantly reduced at the end of the flight training (β = -0.006, p < 0.01; Fig. 4C), indicating that the specific path that birds take becomes more consistent with experience. We also quantified the stereotypy of the horizontal position of the bounds in the flight trajectory. To visualize this, for each bird we plotted the location of the middle of the bound as a heatmap, arranged from the earliest flights (Day 1, first row) to the latest flights (Day 10, last row) then averaged those heatmaps across all birds (Fig. 4B). While most birds performed bounds on Day 1, as the heatmaps illustrate (row 1 of Fig 4B), bound positions were generally spread along the cage length axis and there was substantial variation both within and between individuals in the location of bounds. The bin with the maximal overlap between birds on Day 1 was bin 7 (of 14) which had an overlap of 48%. Beginning around Day 3, this maximum overlap shifted closer to the landing perch (bin 8 of 14) and increased to 67% overlap. The overlap within that location further increased to a maximum of 85% on Day 10. These results show that the experience-dependent increase in bound ratio described earlier (Fig. 2C,E) is accompanied by an increase in the consistency of bound position along the flight horizontal trajectory.

In general, bound positions coincided with a peak in the altitude. Such a relationship aligns with visual observations of small birds in flight, where bounds resemble ’jumps’ in their motion. Consistent with this, we found that the maximum altitude of the beak (the difference between maximum and minimum beak y-coordinates) significantly increased with the time spent bounding (BR) (β = 1.12, p < 0.0001). Bounds tended to begin just before birds reached the altitude peak such that birds bounded during the ascent to and descent from the peak altitude. This was particularly true when birds performed a long, single bound during their flight. The horizontal location of the altitude peak became more stereotyped with experience. When we extracted the altitude peak of the longest bound for each flight, we found that the standard deviation of their horizontal positions was significantly lower on the last day of flight experience compared to the first day (Fig. 4D β = -0.516, p < 0.0001). This finding provides a direct measure of the increased consistency in bound positioning with experience, as reflected by reduced variability in bound peak positions over time.

### Experience-dependent changes to flight show features of motor skill learning

The changes to flight that occur with experience, in particular the increase in speed and the decrease in flight variability, share a number of similarities with other examples of motor skill learning. To test the degree to which the changes to flight that we see with experience are examples of motor practice, we focused on two hallmarks of motor skill learning: faster relearning compared to original learning of flight features (known as ‘savings’) and coordinated increases to speed and accuracy.

First, we investigated the degree to which there were savings of the training-related changes to flight behavior, that is, was the return to peak performance following a pause in training more rapid than the original learning. To this end, once birds had completed 10 days of flight experience, they were returned to group housing for at least two months before being tested again on the perch distance of 70 cm (“retest”). We found that during the retest, birds maintained their flight stereotypy following two months without flying in the corridor. Specifically, the standard deviation of the bound horizontal position was not significantly different on the retest compared to two months prior (Fig 4D β = 0.542, p = 0.338). For flight speed, a comparison of all retest flights found that birds flew significantly faster on the retest compared to Day 1 (t= -2.81 p = 0.0195) but also flew significantly slower on the retest compared to Day 10 (t= 2.76, p = 0.0217). However, they returned to the maximum flight speed by the end of the 20 retest flights (F=7.69; p<0.0001). Specifically, the first five retest flights were significantly slower than those on Day 10 (p=0.0054; Fig. 4E) and not different from Day 1 flights (p=0.1781) while the last five retest flights were significantly faster than Day 1 flights (p=0.0079) and not significantly different from Day 10 flights (p=0.1405).

Second, we examined how accuracy in bound positioning varied with flight speed between the first and last day of practice (Fig 4F). All birds showed a negative slope between speed and the standard deviation of the position of the altitude peak during the bound, indicating that both speed and accuracy improve with experience. Taken together, the simultaneous increases in speed and stereotypy with training and the savings of these parameters after two months without training indicate that flight shows hallmarks of motor practice that we can quantify in our behavioral paradigm.

### Birds flexibly adjust trajectories over increasing distances

Zebra finches adapt their flight behavior to increasing perch distances by boosting their speed and incorporating more short bounds into their trajectories while maintaining a stable wingbeat frequency (See summary of statistical results Table 1). In a subsequent analysis, we examined how the beak trajectories are adjusted to accommodate the greater flight distances. As birds fly over longer distances, one possibility is that they generalize the trajectory that they have consolidated on short flights to longer distances, effectively stretching their flight trajectories and bound positions. Alternatively, they may flexibly adapt their trajectory and bound position depending on the distance. To investigate this, birds that received ten days of flight training performed ten flights in corridors of increasing length (70, 100, 130, 160 cm). As illustrated by the trajectories of one example bird (Fig. 5A), at the shortest distances the trajectories show an inverted-U shape with a single peak. However, on longer flights, the trajectory shape is modified and there is the appearance of a second bound. Across birds, we found similar modulation of the trajectories indicating that birds do not stretch their trajectories to adapt to longer distances but instead adjust the number and location of bounds. When we averaged the bound positions for all birds corresponding to the 70 cm distance along a 1-dimensional heatmap (Fig. 5B), we found a maximal overlap (71%) at the middle, 8th bin, similar to what was observed during the later stages of practice for the 70 cm distance (Fig. 4B). At longer distances, the positions of bounds are more variable between birds, indicated by the less prominent peak in the heatmaps, with maximal values next to the middle of the length axis. The 160 cm heatmap is distinct, showing an initial overlap zone around the middle and a second zone emerging near the landing perch, suggesting a group pattern similar to the individual shown in Fig. 5A.

**Figure 5.**
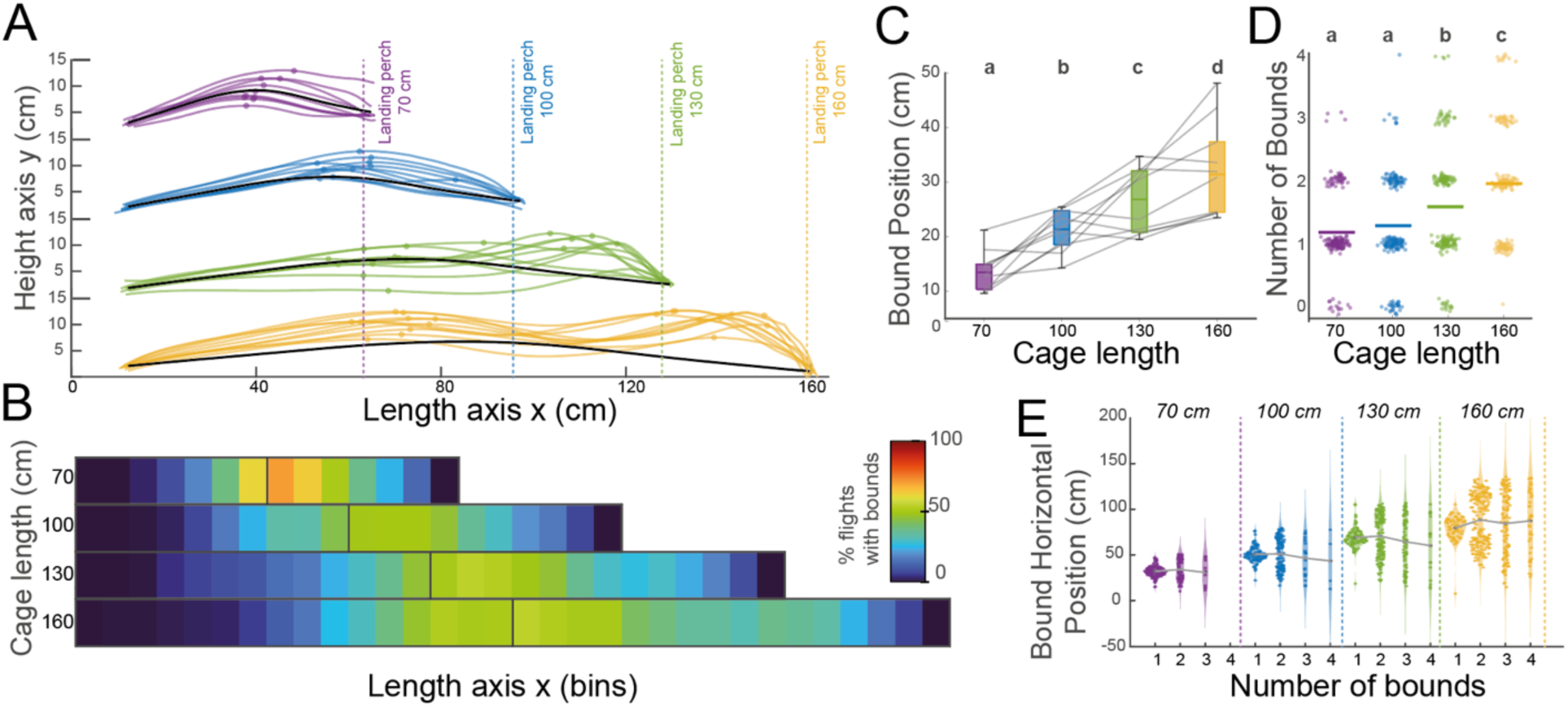
(A) Flight trajectory traces based on beak position for a single bird on 70 cm (purple), 100 cm (blue), 130 cm (green) and 160 cm (yellow) flights. Diamonds indicate mid-bound positions. Black lines are the mean trajectory for the 70 cm flight, linearly stretched to longer distances. (B) Heat map of bound positions for all birds over the four distances from 70 cm (top row) to 160 cm (bottom row). Length axis is divided into bins. Black line are the middle of the length axis. Color indicates the percent of flights across all birds with a bound in that bin (warmer colors are higher percent). (C) Bound positions move progressively farther along the length axis as the flight length increases. Different letters indicate significant differences at p < 0.05. (D) Number of bounds increases with increased flight distance. (E) Plot of all bound positions for each bird at the four flight distances. For each flight distance, the average bound position at that distance (black points and line) is the same regardless of the number of bounds birds perform.

**Table 1:** Summary of statistical results from linear mixed models.

| Protocol | Variable | Covariable | Estimate $\beta$ | p-values |
| --- | --- | --- | --- | --- |
| Practice | Speed | Total rank | 0.002 | < 0.0001 * |
| Practice | Speed | Daily rank | 0.003 | 0.013 * |
| Practice | Bound Ratio | Total rank | 0.363 | < 0.0001 * |
| Practice | Bound Ratio | Daily rank | 0.006 | < 0.0001 * |
| Practice | Beak amplitude | Total rank | 0.276 | < 0.0001 * |
| Practice | Beak amplitude | Daily rank | 1.260 | 0.002 * |
| Practice | Wingbeat freq. | Total rank | -0.000 | 0.240 |
| Practice | Wingbeat freq. | Daily rank | 0.026 | 0.880 |
| Practice | Speed | Bound Ratio | 0.008 | < 0.0001 * |
| Practice | Speed | Wingbeat freq. | -0.001 | 0.692 |
| Practice | Beak amplitude | Bound Ratio | 1.119 | < 0.0001 * |
| Distance | Speed | Distance | 0.557 | < 0.0001 * |
| Distance | Speed | Total rank | 0.007 | < 0.001 * |
| Distance | Bound Ratio | Distance | -4.543 | 0.096 |
| Distance | Bound Ratio | Total rank | 0.169 | 0.032 |
| Distance | n bounds | Distance | 1.051 | < 0.01 * |
| Distance | n bounds | Total rank | -0.014 | 0.167 |
| Distance | Beak amplitude | Distance | 5.0416 | 0.514 |
| Distance | Beak amplitude | Total rank | 0.8636 | < 0.001 * |
| Distance | Wingbeat freq. | Distance | -0.996 | 0.391 |
| Distance | Wingbeat freq. | Total rank | 0.031 | 0.354 |

\* indicates significance at $p < 0.05$

Across all birds, there was a shift in the location of bounds at longer flight distances, with the mid-bound position occurring farther along the corridor at longer distances (Fig. 5C). In addition, while there was substantial variation between birds in the number of bounds produced, there was also an overall increase in the number of bounds (Fig 5D). In particular, while the majority of birds produced a single bound at 70cm, few birds produced a single bound at 160 cm (Fig. 5D). Intriguingly, even though at all distances individual birds produced different numbers of bounds, within a distance, the average middle position of bounds was similar regardless of the number of bounds and occurred roughly at the midpoint of the flight distance (Fig. 5E). Thus, although birds differ in the number of bounds they use at different lengths, they appear to apply similar rules to the positioning of those bounds relative to the middle of the flight distance.

### Learning in single-bound flight is not optimized toward a single goal

To gain further insight into what zebra finches are learning during their bounding flights, we modelled a perch-to-perch flight using two-dimensional kinematic equations of motion over three phases: a flap phase, a bound phase, and a brake phase (Fig. 6A). This model assumes that the zebra finch is a point mass moving under a constant acceleration with no effect of air resistance (see Methods). We modelled flights to test whether females were optimizing towards each of the following objectives: minimizing total flight time, minimizing time spent bounding, maximizing time spent bounding, minimizing velocity at the start of the bound, minimizing deceleration, and minimizing energetic costs. We found that none of these single parameters aligned with the experimental trajectories of single-bound flights. For example, in Fig. 6B the optimized trajectories for each objective (red lines) align poorly with the experimental trajectories (black lines). We also compared the bound locations generated by optimizing for each objective with the bound coordinates for all experimental flights. Again, there was no single optimized objective that aligned with the bound positions from the real data (Fig. 6C). Rather, the experimental bound locations fall into the middle of the optimized values for the different goals (Fig 6C).

**Figure 6.**
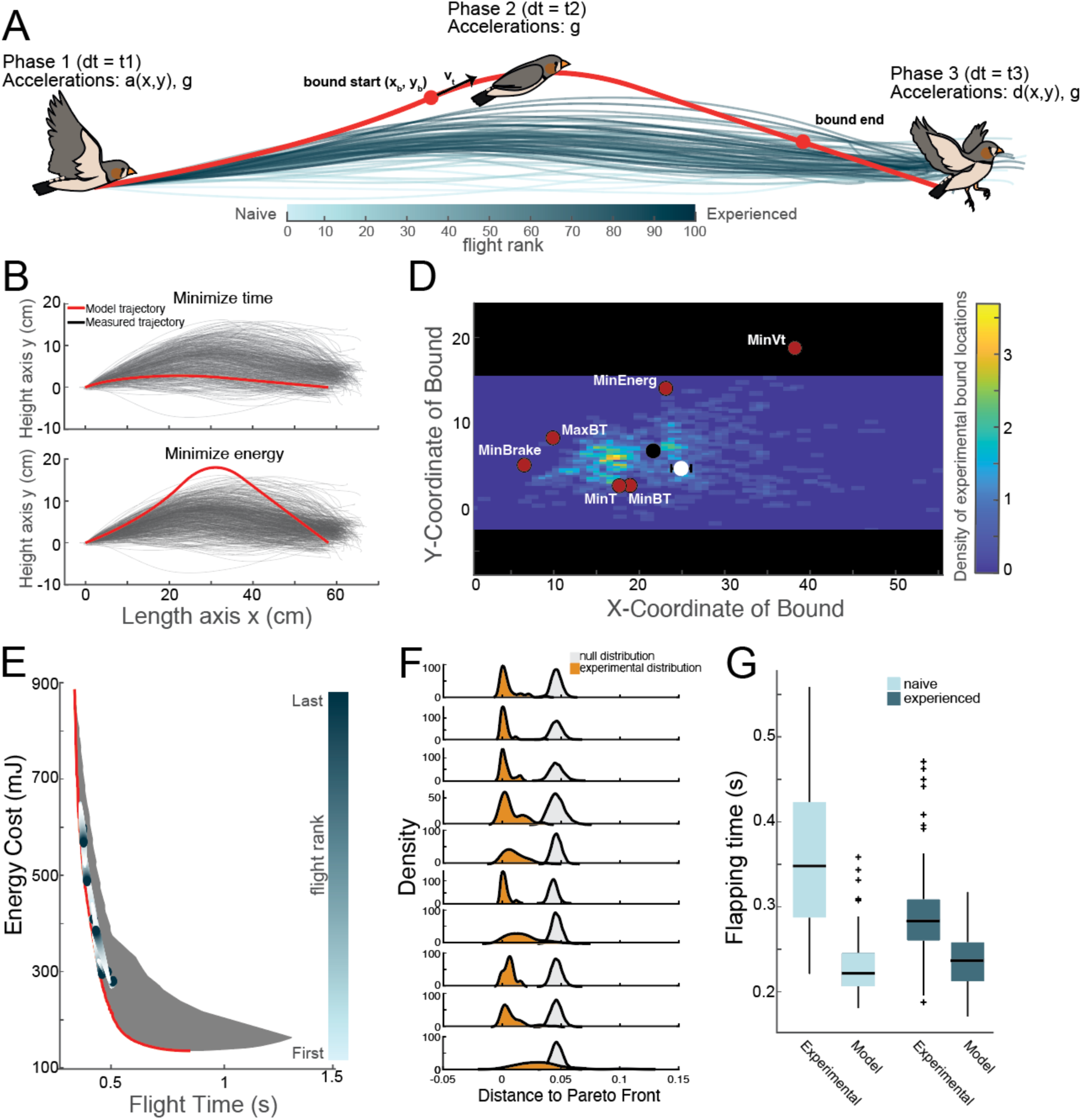
Optimized flight trajectories. (A) Illustration of the model phases and parameters. (B) Model trajectories (red lines) optimized for the minimum time (top) and minimum energy (bottom). Black lines are measured flight trajectories of birds. (C) A density heatmap of the bound coordinates for all experimental flights. The experimental bound locations fall in the middle of optimised values for different goals (shown in red). Also plotted are the mean and standard errors of the bound coordinates for the first (white circle) and last (black circle)15% of flights. (E) Most birds learn bounding positions that shift towards lowering energetic costs while maintaining little increase in flight time. Bounding positions for all birds plotted as a color gradient from the first 15% of flights (light colors) to the last 15% of flights (darker colors). Gray is the entire set of feasible solutions for the analytical model and the pareto frontier is outlined in red. (F) For each bird (rows), experimental trajectories (orange) lie significantly closer in Euclidean distance to the pareto front than random distributions (gray). (G) Experimental trajectories employ longer times spent flapping for a given bounding position compared to model predictions during the first 15% of flights (naïve). These differences are reduced during the last 15% of flights (experienced).

While no single optimization parameter aligned to either the trajectories or bound positions of the experimental flights, the solutions highlighted the possibility that birds may be simultaneously optimizing for more than one objective. To investigate this, we generated the entire set of feasible solutions for analytical models that jointly optimized two objectives. In particular, we focused on the objectives of minimising energetic costs and flight time as experimental trajectories lay between trajectories with those two objectives (Fig 6B). We found that bounding positions sit at the pareto frontier for a model that optimizes to minimize energetic costs and flight time (Fig 6E). For all birds, the experimental trajectories lie closer to the pareto front than random (Fig 6F). Intriguingly, over the course of training, birds generally learn bounding positions that follow the pareto front, shifting towards lowering energetic costs with minimal increases in flight time (Fig 6E). Taken together, these data indicate that with motor practice birds adjust the position and consistency of bounds, thereby potentially decreasing their energetic costs without greatly increasing flight time.

The models predicted that birds would show a slight increase in flight duration with learning while, in contrast, the duration of actual flights decreased with training (Fig. 2B). To understand what could contribute to this difference, we compared the flights of birds and the predicted flights from the model equations performing bounds at similar locations. We found that birds spent longer in the flapping phase for a given bound initiation position on early training flights than the models predicted (Fig. 6G). However, at the end of training, the flight durations are similar between the bird flights and the models. As our models were constrained by an assumption of constant acceleration during this flapping phase, it is possible that these trajectories with longer flapping phases would require time-varying acceleration profiles to achieve the same bound initiation position and velocity to make successfully arrive at the landing perch. As such, these differences raise the possibility that the models may underestimate the energetic savings that occur with training by not being able to account for all possible trajectories.

## DISCUSSION

Birds are capable of performing elaborate flight maneuvers in variable environmental conditions. While it is clear that flight is an adaptable and skilled motor behavior, we know surprisingly little about how birds are able to master this ability. Across species, skilled motor behaviors show practice-related changes or improvements in performance^3^. Understanding which features of a motor behavior change with learning can lend insight into the constraints, flexibility, and optimization of motor behavior. Here, we combined high-speed video recordings and pose-tracking software to analyze the kinematics of thousands of flap-bounding flights in zebra finches over multiple days of flight training and over different distances. We found that birds increase their speed and the time spent bounding and reduce variability in the bound position over ten days of flight performance. These motor changes to flight show savings, as performance is maintained even after two months without flight experience. Moreover, these same parameters are adjusted when birds fly longer distances, indicating that they may be key to flight flexibility. Taken together, our data highlight that flap-bounding flight shows hallmarks of skilled motor learning and lend new insight into the function of bounds.

Motor skill learning refers to the process by which movements are executed more quickly and accurately with practice. Such learned movements are easier to relearn than to learn from scratch, a phenomenon known as ‘savings’ that is often used as a litmus test of memory consolidation^3,19–21^. Flight exhibits both of these hallmarks of motor practice. Following ten days of flight experience, birds flew faster with more consistent trajectories and repeatable bound positions and birds improved both parameters simultaneously over the course of practice. In addition, when birds were retested on the same 70 cm distance following at least two months without training, they rapidly returned to performing flights with similar stereotypy of the bound position and at the higher speeds seen on the last day of training, indicating savings of the learned motor pattern. While savings are often associated with motor consolidation, future investigation of whether there is a period during which the motor memory is sensitive to interference before becoming stabilized will provide further support for the degree to which there is consolidation of flight motor memories.

Birds adjusted similar flight parameters over the course of motor practice and across different flight distances, including the location and duration of bounds, the flight speed, and the consistency of bound locations and trajectories. In contrast, they did not appear to adjust their flapping rate either across days of training or over different distances. Indeed, the flapping rate was highly stable across the 10 days of training and from 0.7 to 1.6 meters. These data fit with previous work that has found that birds are more likely to modulate their wingbeat amplitude than their wingbeat frequency during more energy-demanding flight modes^22^ and that wingbeat frequency has been shown to relate to breathing rate^23–25^. Taken together, these data highlight that bounds, rather than flapping, may be a key node for adjustment and flexibility in intermittent flight.

Previous work has developed models to explain why small birds include bounds in their flight, especially at lower speeds or shorter flight distances. For example, one longstanding hypothesis for bounding, the “fixed-gear” model, proposed that small birds are constrained in motor-unit recruitment^18^. The model therefore predicted that whenever less-than-maximum power is required, small birds should use intermittent bounds. More recent data found that zebra finches show substantial variation in motor-unit recruitment, indicating that birds are not bounding because they are constrained to a small range of power^26,27^. However, the function of bounds in short flights remains unclear. We used two-dimensional kinematic equations to model flight towards multiple different objectives and found that birds stay near trajectories that simultaneously optimize energy cost and flight time. Birds are not only closer to the pareto front than random, but birds move along the pareto front over the course of practice, decreasing energy with only slight changes to flight time. These data hint that even at slow flight speeds over short distances birds may adjust bounds and trajectories to optimize energy and time costs of flight.

Intriguingly, comparison of our models to the actual flights found that birds spent longer in the flapping phase and flew at slower speeds early in practice than what the models predicted. While our models were constrained to a constant acceleration during the flapping phase, birds were not as constrained, resulting in the mismatch in flapping time especially during their initial flights. However, in later flights, as the probability of bounding as well as the bound location become more consistent, there is better coherence between the data and the model suggesting that model assumptions are more applicable to model these behaviours. A shift towards a more constant acceleration during flapping with experience is perhaps unsurprising as studies on pigeons have shown a similar reduction in kinematic variability (velocity) during landing with familiarity^28^. As our model does not include time-varying acceleration profiles for the flapping phase, it is likely that there are possible trajectories not included in the analysis and that it may underestimate the possible reduction in energy costs associated with a given bounding profile. Further investigation of the kinematics of the wings during the flapping phase, in particular quantifying whether there is variation in the amplitude or rhythm of flapping, as well as looking in greater detail at changes in acceleration during flapping will lend further insight into the optimization of flap-bounding flight.

Practice induces learning-dependent changes in functional brain networks and these are thought to represent a motor memory^29–32^. While practice-related neural changes and motor memory encoding are the focus of considerable research in mammals, investigation of the neural basis for sensorimotor practice in other species has focused on a limited suite of behaviors in a subset of animals. For example, in songbirds, the circuits for learning and performance of vocal-motor behavior are well-studied^33,34^, however, there has been little investigation of circuitry for learning and flexibility of other motor behaviors. Moreover, studies on neural circuits for flight have focused on sensory inputs including somatosensory responses and visual motion^35–39^. Future work delineating the neural circuits for flexibly modulating flight will lend new insight into shared principles of sensorimotor learning and integration across species.

## METHODS

### Animals

Ten female Zebra finches (*Taeniopygia castanotis*) were used in this study (age > 90 days post-hatch, average weight 14.4± 1.4 g). Birds were raised in a breeding colony (McGill University) with parents and siblings until 60 days of age then housed in same-sex group cages (dimensions 40 x 40 x 35 cm). Birds were maintained on a 14/10 light-dark cycle with ad libitum access to seed, water, and grit, as well as vegetables and egg supplements once per week. Animal care and procedures followed all Canadian Council on Animal Care guidelines and were approved by the Animal Care Committee of McGill University.

### Flight corridor

All flight experiments were performed in a transparent plexiglass corridor (CPI Automation, Ontario, Canada, Fig. 1A) 2 m × 0.3 m × 0.6 m (length × width × height) lit by 4 non-flickering LED panels over the top (Genaray Spectro LED Essential 500IID). Within the corridor, one intermediate plexiglass panel could slide along the length axis to vary the available distance for flight. For each session, we placed a piece of paper, folded in half to create a peak, onto the floor of the corridor to prevent birds from landing on the floor. Two 8 cm wooden perches, one at each end of the corridor, were affixed 40 cm above the floor. Each perch was connected to a micro switch (GPTCLS01, ZF Friedrichshafen AG) to detect perch movements (e.g. take-offs and landings). The binary signal coming from switches is sent to a laptop via an input/outputdevice (National instrument NIDAQ 6501) and landing and takeoff time points were extracted using custom written software (Matlab, Mathworks, Natick, MA).

### Behavioral protocols

Birds were motivated to perform back and forth flights with food and social rewards. Food rewards consisted of a small amount of crushed, boiled egg in a dish next to one of the perches. For social rewards, a rectangular piece of Smart Glass (Smart Glass Technologies, Toronto Ontario) would turn on (become transparent) when birds landed on a perch connected to a microswitch positioned in front of the glass. Once birds landed on the perch, the smart glass would become transparent for 5 seconds or until the bird took off for another flight, whichever happened first. While the glass is transparent, the focal bird can see a second female from the experimental group, randomly chosen at the beginning of the session and placed in a small cage in the other side of the SmartGlass. In case the bird was still reluctant to fly the experimenter waved a hand close to the bird to initiate takeoff.

Prior to the experiments, zebra finches were introduced to the corridor as a group (n=2-5 birds/group) to acclimate to the chamber. This habituation session ended when each bird was able to fly between the two perches, land, and perch without dropping to the floor. Birds were then tested in two experiments. First, we investigated the effects of flight experience on flight parameters (“flight training experiment”). Zebra finches’ flight behavior was evaluated during 10 consecutive days with one session per bird and per day. Each session started by placing the focal bird in the flight arena (70cm end-to-end; 56 cm from beak tip to landing perch) and another female behind the SmartGlass and ended when 20 back and forth flights were recorded. Session duration was dependent on the bird’s motivation and varied between 2 to 55 minutes (mean=13.9). We next evaluated the effect of distance on flight behavior (“flight distance experiment”) beginning two months after the completion of flight training experiments. For this experiment, we gradually increased the flight distance by sliding the intermediate panel in the corridor, extending it from 70 cm to 100 cm, 130 cm, and finally 160 cm. When the bird made 20 back and forth flights at one distance the session ended and she was tested at the next distance the following day. Session duration varied from 5 to 36 minutes (mean=14.9). For both protocols, flights in which the bird missed the landing perch, touched the floor, or tried to perch elsewhere were discarded and not part of the analysis.

### Video processing and DeepLabCut tracking

Videos were acquired with a GoPro camera (Gopro10 Black, Narrow view, 1080p, 240fps) mounted on 80/20 slots. The camera was positioned so that the entire length of the corridor was in frame. The camera was fixed at a distance of 14 cm from the corridor for flight training experiments and 122 cm from the corridor for flight distance experiments. We used the landing and takeoff timestamps from the perch microswitches to cut the full, raw videos into multiple, single flight videos grouped by flight direction: flying from the left to the right perch (*left2right*) or from the right to the left perch (*right2left*). All cropped videos were visually checked and those where the bird touched the side or bottom of the corridor, or where the camera field of view was obstructed were discarded from the analysis. We accumulated videos of a total of 2,009 single flights for dataset 1 (left2right: 1014; right2left: 995) and 800 for dataset 2 (left2right: 400; right2left: 400) (Supplementary table 1).

For all single-flight videos, we computed the transformation matrix (*imregtform*) of each frame towards a reference frame to account for small differences in camera position across videos. Single-flight videos were then input to the DeepLabCut (DLC) framework^40,41^. We manually labeled the position of the bird’s beak, tail-tip, wing tips and the two perches in a subset of the videos (Fig 1B). For flight training experiments, both wingtips were labelled. For flight distance experiments, because of the increased camera distance needed to have the entire corridor in frame, only one wingtip could be reliably identified and labeled. In total, 4 separate DLC projects were created for the two protocols and flight directions (Supplementary table 1 for details). Each project is trained on a distinct set of videos designed to minimize variability. For example, the ’practice Left2Right ’ project includes videos of all birds flying from left to right, recorded according to the configuration of the flight training experiment. One output file was generated per video after analysis with x and y coordinates of all body parts at every frame. When wing-tip tracking results were discontinuous or had low likelihood values, additional videos were added to the training set, initiating another iteration of refinement. All frames with a likelihood below 0.6 were excluded in final analysis.

To account for slight variations in camera positions across videos, we estimated the geometric transformation needed to align each frame to a common reference frame used for all videos in the DLC project (using *imregtform* in Matlab). However, this initial alignment does not fully correct for differences in perch positions between flight occurrences. To address this, we computed a second transformation based on aligning the two perch positions to their corresponding locations in the reference frame (using *fitgeotform2d* in Matlab). These two consecutive transformations were then applied to the coordinate files obtained with DLC.

To gain further precision about take-off and landing times, we leveraged the fact that birds induce a vertical movement of the perch when they take off and land. We tracked perch position and used changes in the y-coordinates to identify landing and takeoff times for each flight. Only the frames between these events were included in the analysis.

### Kinematic analysis

We first divided flight into flapping and bounding components. Flapping is defined as periods with up and down motion of the wing tips while bounds are periods without flapping when birds bring their flexed wings in close to the body (Fig. 1C). Using the registered coordinates of the wing tips, we identified periods of flapping and bounding. For this, we generated a line of the body axis by connecting the beak and tail coordinates, then calculated wing elevation as the shortest distance of the wing tip from the beak-tail tip line at every frame (Fig. 1B). A positive or negative elevation reflected an up or down wing stroke respectively. Elevation was calculated for the wing closest to the camera (right wing when birds flew left to right and the left wing when birds flew right to left). For bound detection, we computed the local standard deviation (*stdfilt*) of the wing elevation over time. We defined a bound as a phase of very low deviation across at least 5 consecutive frames and extracted the central position and duration in frames for each bound for analysis.

For each flight, we quantified various features including the speed, percent of time bounding, wingbeat frequency, and beak amplitude. Flight speed (m.s^-1^) was calculated as: *distp/((fland-ftakeoff)/fr)* where distp=distance between perches in meters; f=frame; fr=framerate in frames per seconds. The ratio of bounds in the flight (BR) was defined as: number of bound frames / total number of frames of the flight * 100. Wingbeat frequency (Hz) represents the number of wingbeats per duration of a flapping phase (in s) / fr. Beak amplitude (pixels) was calculated as the difference between the beak maximal and minimal y coordinate of the flight trajectory.

We tested the effect of practice by modelling kinematics as a proportion of flight rank using linear mixed effects models (*fitlme* function, MATLAB 2022b). Total flight speed, BR, beak amplitude and wingbeat frequency each represented a dependant variable of an LME model. We entered total flight rank, daily rank and direction of flight (n=2) as fixed effects while taking into account bird ID (n=10) as a random effect. The final formula of the test was: *‘KinematicParam ∼ totalrank + dayrank + direction + (1 |ID)’*. In protocol 2, birds always started with the smallest distance between perches. Hence, in order to test the effect of distance, we entered cage length parameter (n=4) but also total flight rank and direction as fixed effects of the same kinematic data, with bird ID as random effect using the formula: ‘*KinematicParam ’ ∼ cagelength + totalrank + direction + (1 |ID)*’. Interactions between two kinematic variables were tested with the following LME’s formula: *‘Param1 ∼ Param2 + direction + (1 |ID)’*. The number of bounds was processed differently since it is a count variable. We tested its relation with flight rank and distance using the same fixed and random effects but with a generalized mixed model with a Poisson distribution (*fitglme* function, MATLAB 2022b).

Flight trajectories were computed from the x,y beak coordinates obtained with DeepLabcut and filtered at 0.6 likelihood minimum. They were trimmed between the landing (fland) and takeoff frames (ftakeoff) then aligned to the takeoff perch position and finally smoothed using a 3 degree-Savitzky-Golay filter (matlab smoothdata). Comparison of perch spacing in the cage versus on the images allowed us to convert coordinates from pixels to centimeters. Trajectories were interpolated to ensure a consistent number of points, then compared two-by-two using a pairwise Euclidian distance with the following formula: sqrt(sum((x1 − x2). ^2 + (y1 − y2). ^2)). All distances were averaged per day and across birds to test their variation with practice, using a right-tailed Wilcoxon signed-rank test.

We employed a heatmap-based method to visualize the effect of practice on bound positioning more easily, and average the data across bird IDs. Heatmaps of bounds position were generated from histogram-counts of their x, y coordinates into bins of equal size. Values of the y-axis were summed to obtain one-dimensional heatmaps along the cage length axis that were then binarized for each flight. Finally, values were divided by the total number of flights and multiplied by 100 to obtain the proportion of flights containing a bound at each bin location. Heatmaps of the right2left direction were flipped horizontally for visualization purpose. Average heatmaps were generated across bird IDs and the two flight directions.

### Modeling of the optimal trajectories

#### Trajectory Optimisation

##### 1. Model Equations

To test hypotheses on what the zebra finches are learning during their bounding flights, we modelled a perch-to-perch flight using two-dimensional kinematic equations of motion over three phases: a flap phase, a bound phase, and a brake phase. This model assumes that the zebra finch is a point mass moving under a constant acceleration with no effect of air resistance.

The flap phase was defined by:

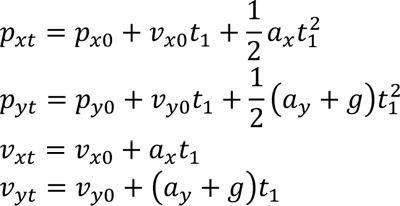

where 𝑝*_xo_*, 𝑝*_yo_* are the starting coordinates of the flap phase, 𝑝*_xt_*, 𝑝*_yt_* are the ending coordinates of the flap phase, 𝑣*_x_*_0_, 𝑣*_y_*_0_ are the velocities at the start of the flap phase, 𝑣*_xt_*, 𝑣*_yt_* are the velocities as the flap phase transitions to the bound phase, 𝑎_x_, 𝑎_y_ are the accelerations due to flapping, 𝑔 is gravitational acceleration, and 𝑡_1_ is the duration of the flap phase.

The bound phase was defined by:

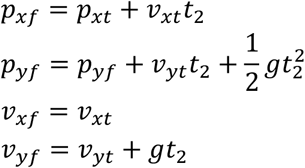

where 𝑝*_xf_*, 𝑝*_xf_* are the ending coordinates of the bound phase, 𝑣*_xf_*, 𝑣*_yf_* are the velocities at the end of the bound phase, and 𝑡_2_ is the duration of the bound phase.

The brake phase was defined by:

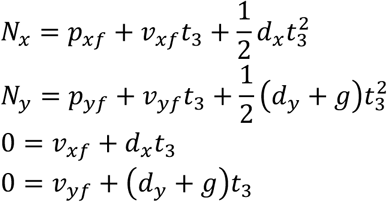

where 𝑁_*x*_, 𝑁_*y*_ are the ending coordinates of the brake phase, 𝑑*_x_*, 𝑑*_y_* are the accelerations due to braking, and 𝑡_3_ is the duration of the brake phase.

##### 2. Optimisation Methodology

To solve this system of equations, we assigned the following variables known values:

| Variable | Value |
| --- | --- |
| $p_{x0}$ | 0 m |
| $p_{y0}$ | 0 m |
| $v_{x0}$ | 1.54 m/s* |
| $v_{y0}$ | 0.6 m/s* |
| $N_x$ | 0.58 m** |
| $N_y$ | 0 m |
| $g$ | -9.81 m/s<sup>2</sup> |

\* average value derived from experimental data
\*\* cage length derived from experimental flight chamber

In addition to this, we have three optimisation variables:

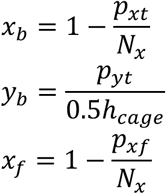

where ℎ_*cage*_ = 0.46𝑚, 𝑥_*b*_𝜖[0.1,0.9], 𝑦_*b*_𝜖[0.1,1], and 𝑥_*f*_𝜖[0.1,0.9]. This results in twelve unknowns, resulting in a unique solution to these equations. However, we further impose a set of constraints to generate only realistic solutions:

a. Time must be positive: 𝑡_1_ ≥ 0, 𝑡_2_ ≥ 0, 𝑡_3_ ≥ 0
b. Flapping induced accelerations must not exceed a critical value: 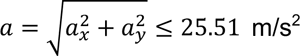 and 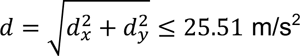 (pied flycatchers had a maximum of g-force of 2.6g during upwards escapes^42^)
c. _(c)_ The maximum height cannot exceed the experimental flight chamber: ℎ*_max_* ≤ 0.5ℎ*_cage_*
d. The bound length must exceed a minimum value, preventing the solver from trying to remove the bound completely by putting the start and end of the bound infinitely close to one another: 𝑝*_xf_* – 𝑝*_xf_* ≥ 0.14m. This value was derived from an average of the single bound flights from experimental data.

We can then solve the problem using the particle swarm algorithm in Python’s *pyswarm* library in order to minimise either of the following objectives:

A. The birds are learning to minimise their velocity at the start of the bound: min 𝑣*_t_* where 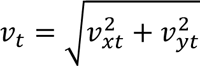
B. _B._ The birds are learning to minimise their total flight time: min 𝑇 where 𝑇 = 𝑡_1_ + 𝑡_2_ + 𝑡_3_
C. _C._ The birds are learning to minimise time spent during a bound, when there’s less control authority: min 𝑡_2_
D. _D._ The birds are learning to maximise time spent during a bound: max 𝑡_2_
E. The birds are learning to minimise their deceleration or brake rate: min 𝑑
F. The birds are learning to minimise their energetic costs: min 𝐸 where 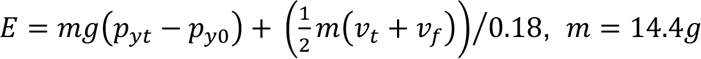, and 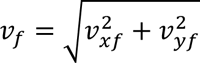. We divided the kinetic energy portion by 0.18 because the efficiency of flight muscles in small birds is estimated to be between 13%-23%^43,44^

---

**Supplementary Table 1:** DeepLabCut settings.

| Projects | Protocol 1 Practice<br>Left2right direction | Protocol 1 Practice<br>Right2left direction | Protocol 2 Distance<br>Left2right direction | Protocol 2 Distance<br>Right2left direction |
| --- | --- | --- | --- | --- |
| Training set size<br>Videos (frames) | 28 (833) | 28 (870) | 20 (494) | 23 (638) |
| Number of iterations | 2 | 2 | 1 | 2 |
| Network | Resnet_50 | Resnet_50 | Resnet_50 | Resnet_50 |
| Network iterations | 500k | 300k | 400k | 350k |
| Train error (px) | 2.8 | 3.82 | 3.63 | 1.59 |
| Test error (px) | 7.02 | 9.83 | 2.87 | 2.82 |
| Final number of videos<br>(flights) | 1014 | 995 | 400 | 400 |

## ACKNOWLEDGEMENTS

We would like to thank David Perkel for helpful discussions and comments on the manuscript. In addition, we would like to thank Katherine Chadwick and Sophie Morris for assistance with filming, and Myriah Haggard for help with data curation. This work was funded by the Human Frontiers in Science Program to S.C.W. and S.W. (RGP0068/2021) and the Natural Sciences and Engineering Research Council of Canada (NSERC) to S.C.W.

## AUTHOR CONTRIBUTIONS

C.B. and S.C.W. designed the experiments; C.B. and S.C.W. performed the experiments; J.W. and S.W. created optimized models, C.B., J.W., and S.C.W. analyzed the data; and C.B., J.W., and S.C.W. wrote and C.B., J.W., S.W., and S.C.W. edited the manuscript.

## DECLARATION OF INTERESTS

The authors declare no competing interests.

## Notes

### Competing Interest Statement

The authors have declared no competing interest.

## REFERENCES

1. Shmuelof, L., and Krakauer, J.W. (2011). Are We Ready for a Natural History of Motor Learning? Neuron 72, 469–476. 10.1016/j.neuron.2011.10.017.

2. Dhawale, A.K., Smith, M.A., and Ölveczky, B.P. (2017). The Role of Variability in Motor Learning. Annual Review of Neuroscience 40, 479–498. 10.1146/annurev-neuro-072116-031548.

3. Motor Learning - Krakauer - 2019 - Comprehensive Physiology - Wiley Online Library.

4. Woolley, S.C., and Kao, M.H. (2015). Variability in action: Contributions of a songbird cortical-basal ganglia circuit to vocal motor learning and control. Neuroscience 296, 39–47. 10.1016/j.neuroscience.2014.10.010.

5. Mooney, R. (2009). Neurobiology of song learning. Current Opinion in Neurobiology 19, 654–660. 10.1016/j.conb.2009.10.004.

6. Brainard, M.S., and Doupe, A.J. (2002). What songbirds teach us about learning. Nature 417, 351–358. 10.1038/417351a.

7. Derégnaucourt, S., Mitra, P.P., Fehér, O., Pytte, C., and Tchernichovski, O. (2005). How sleep affects the developmental learning of bird song. Nature 433, 710–716. 10.1038/nature03275.

8. Mizuguchi, D., Sánchez-Valpuesta, M., Kim, Y., dos Santos, E.B., Kang, H., Mori, C., Wada, K., and Kojima, S. (2024). Daily singing of adult songbirds functions to maintain song performance independently of auditory feedback and age. Commun Biol 7, 598. 10.1038/s42003-024-06311-5.

9. Whishaw, I.Q., and Pellis, S.M. (1990). The structure of skilled forelimb reaching in the rat: A proximally driven movement with a single distal rotatory component. Behavioural Brain Research 41, 49–59. 10.1016/0166-4328(90)90053-H.

10. Buitrago, M.M., Ringer, T., Schulz, J.B., Dichgans, J., and Luft, A.R. (2004). Characterization of motor skill and instrumental learning time scales in a skilled reaching task in rat. Behavioural Brain Research 155, 249–256. 10.1016/j.bbr.2004.04.025.

11. Klein, A., and Dunnett, S.B. (2012). Analysis of Skilled Forelimb Movement in Rats: The Single Pellet Reaching Test and Staircase Test. Current Protocols in Neuroscience 58, 8.28.1-8.28.15. 10.1002/0471142301.ns0828s58.

12. Tobalske, B.W., Peacock, W.L., and Dial, K.P. (1999). Kinematics of flap-bounding flight in the zebra finch over a wide range of speeds. Journal of Experimental Biology 202, 1725–1739. 10.1242/jeb.202.13.1725.

13. Schiffner, I., Vo, H.D., Bhagavatula, P.S., and Srinivasan, M.V. (2014). Minding the gap: in-flight body awareness in birds. Front Zool 11, 64. 10.1186/s12983-014-0064-y.

14. Schiffner, I., Perez, T., and Srinivasan, M.V. (2016). Strategies for Pre-Emptive Mid-Air Collision Avoidance in Budgerigars. PLOS ONE 11, e0162435. 10.1371/journal.pone.0162435.

15. Usherwood, J.R. (2016). Physiological, aerodynamic and geometric constraints of flapping account for bird gaits, and bounding and flap-gliding flight strategies. Journal of Theoretical Biology 408, 42–52. 10.1016/j.jtbi.2016.07.003.

16. Duerr, A.E., Miller, T.A., Lanzone, M., Brandes, D., Cooper, J., O’Malley, K., Maisonneuve, C., Tremblay, J.A., and Katzner, T. (2015). Flight response of slope-soaring birds to seasonal variation in thermal generation. Functional Ecology 29, 779–790. 10.1111/1365-2435.12381.

17. Warrick, D.R., Tobalske, B.W., and Powers, D.R. (2005). Aerodynamics of the hovering hummingbird. Nature 435, 1094–1097. 10.1038/nature03647.

18. Rayner, J.M.V. (1985). Bounding and undulating flight in birds. Journal of Theoretical Biology 117, 47–77. 10.1016/S0022-5193(85)80164-8.

19. Malone, L.A., Vasudevan, E.V.L., and Bastian, A.J. (2011). Motor Adaptation Training for Faster Relearning. J. Neurosci. 31, 15136–15143. 10.1523/JNEUROSCI.1367-11.2011.

20. Roemmich, R.T., and Bastian, A.J. (2015). Two ways to save a newly learned motor pattern. Journal of Neurophysiology 113, 3519–3530. 10.1152/jn.00965.2014.

21. Krakauer, J.W., Ghez, C., and Ghilardi, M.F. (2005). Adaptation to Visuomotor Transformations: Consolidation, Interference, and Forgetting. J. Neurosci. 25, 473–478. 10.1523/JNEUROSCI.4218-04.2005.

22. Krishnan, K., Garde, B., Bennison, A., Cole, N.C., Cole, E.-L., Darby, J., Elliott, K.H., Fell, A., Gómez-Laich, A., de Grissac, S., et al. (2022). The role of wingbeat frequency and amplitude in flight power. Journal of The Royal Society Interface 19, 20220168. 10.1098/rsif.2022.0168.

23. Berger, M., Roy, O.Z., and Hart, J.S. (1970). The co-ordination between respiration and wing beats in birds. Z. Vergl. Physiol. 66, 190–200. 10.1007/BF00297778.

24. Butler, P.J., and Woakes, A.J. (1980). Heart Rate, Respiratory Frequency and Wing Beat Frequency of Free Flying Barnacle Geese Branta Leucopsis. J Exp Biol 85, 213–226. 10.1242/jeb.85.1.213.

25. Funk, G.D., Sholomenko, G.N., Valenzuela, I.J., Steeves, J.D., and Milsom, W.K. (1993). Coordination of Wing Beat and Respiration in Canada Geese During Free Flight. J Exp Biol 175, 317–323. 10.1242/jeb.175.1.317.

26. Tobalske, B.W., Puccinelli, L.A., and Sheridan, D.C. (2005). Contractile activity of the pectoralis in the zebra finch according to mode and velocity of flap-bounding flight. Journal of Experimental Biology 208, 2895–2901. 10.1242/jeb.01734.

27. Tobalske, B.W. (2016). Evolution of avian flight: muscles and constraints on performance. Philosophical Transactions of the Royal Society B: Biological Sciences 371, 20150383. 10.1098/rstb.2015.0383.

28. Green, P.R., and Cheng, P. (1998). Variation in Kinematics and Dynamics of the Landing Flights of Pigeons on a Novel Perch. J Exp Biol 201, 3309–3316. 10.1242/jeb.201.24.3309.

29. Kami, A., Meyer, G., Jezzard, P., Adams, M.M., Turner, R., and Ungerleider, L.G. (1995). Functional MRI evidence for adult motor cortex plasticity during motor skill learning. Nature 377, 155–158. 10.1038/377155a0.

30. Kantak, S.S., and Winstein, C.J. (2012). Learning–performance distinction and memory processes for motor skills: A focused review and perspective. Behavioural Brain Research 228, 219–231. 10.1016/j.bbr.2011.11.028.

31. Steele, C.J., and Penhune, V.B. (2010). Specific Increases within Global Decreases: A Functional Magnetic Resonance Imaging Investigation of Five Days of Motor Sequence Learning. J. Neurosci. 30, 8332–8341. 10.1523/JNEUROSCI.5569-09.2010.

32. Pascual-Leone, A., Grafman, J., and Hallett, M. (1994). Modulation of Cortical Motor Output Maps During Development of Implicit and Explicit Knowledge. Science 263, 1287–1289. 10.1126/science.8122113.

33. Brainard, M.S., and Doupe, A.J. (2002). What songbirds teach us about learning. Nature 417, 351–358. 10.1038/417351a.

34. Mooney, R. (2009). Neurobiology of song learning. Current Opinion in Neurobiology 19, 654–660. 10.1016/j.conb.2009.10.004.

35. Gaede, A.H., Baliga, V.B., Smyth, G., Gutiérrez-Ibáñez, C., Altshuler, D.L., and Wylie, D.R. (2022). Response properties of optic flow neurons in the accessory optic system of hummingbirds versus zebra finches and pigeons. Journal of Neurophysiology 127, 130–144. 10.1152/jn.00437.2021.

36. Smyth, G., Baliga, V.B., Gaede, A.H., Wylie, D.R., and Altshuler, D.L. (2022). Specializations in optic flow encoding in the pretectum of hummingbirds and zebra finches. Current Biology 32, 2772–2779.e4. 10.1016/j.cub.2022.04.076.

37. Wylie, D.R., Gutiérrez-Ibáñez, C., Gaede, A.H., Altshuler, D.L., and Iwaniuk, A.N. (2018). Visual-Cerebellar Pathways and Their Roles in the Control of Avian Flight. Frontiers in Neuroscience 12.

38. Gaede, A.H., Wu, P.-H., and Leitch, D.B. (2024). Variations in touch representation in the hummingbird and zebra finch forebrain. Current Biology 34, 2739–2747.e3. 10.1016/j.cub.2024.04.081.

39. Gutiérrez-Ibáñez, C., Wylie, D.R., and Altshuler, D.L. (2023). From the eye to the wing: neural circuits for transforming optic flow into motor output in avian flight. J Comp Physiol A 209, 839–854. 10.1007/s00359-023-01663-5.

40. Mathis, A., Mamidanna, P., Cury, K.M., Abe, T., Murthy, V.N., Mathis, M.W., and Bethge, M. (2018). DeepLabCut: markerless pose estimation of user-defined body parts with deep learning. Nature Neuroscience 21, 1281–1289. 10.1038/s41593-018-0209-y.

41. Nath, T., Mathis, A., Chen, A.C., Patel, A., Bethge, M., and Mathis, M.W. (2019). Using DeepLabCut for 3D markerless pose estimation across species and behaviors. Nature Protocols 14, 2152–2176. 10.1038/s41596-019-0176-0.

42. Tomotani, B.M., and Muijres, F.T. (2019). A songbird compensates for wing molt during escape flights by reducing the molt gap and increasing angle of attack. J Exp Biol 222, jeb195396. 10.1242/jeb.195396.

43. Ward, S., Möller, U., Rayner, J.M.V., Jackson, D.M., Bilo, D., Nachtigall, W., and Speakman, J.R. (2001). Metabolic power, mechanical power and efficiency during wind tunnel flight by the European starling Sturnus vulgaris. J Exp Biol 204, 3311–3322. 10.1242/jeb.204.19.3311.

44. Askew, G.N., and Ellerby, D.J. (2007). The mechanical power requirements of avian flight. Biol Lett 3, 445–448. 10.1098/rsbl.2007.0182.

